# A ribosomal kinetic checkpoint governs selective mRNA recruitment

**DOI:** 10.64898/2026.06.12.731988

**Authors:** Masaaki Sokabe, Carlos Alvarado, Christopher P. Lapointe, Nancy Villa, Joseph D. Puglisi, Christopher S. Fraser

## Abstract

Translation initiation begins with recruitment of mRNA to the ribosome, yet how mRNA engagement is converted into productive initiation remains unclear. Using real-time fluorescence assays with purified components, we show that mRNA recruitment proceeds through a branched kinetic pathway on the 40S subunit. Following rapid sampling, mRNAs partition into either a productive accommodated state or an arrested state that stabilizes ribosome binding before accommodation. mRNA structure, eIF3, and eIF3j bias recruitment toward arrest, whereas eIF4F promotes accommodation in an ATP-dependent manner coupled to displacement of eIF3j from the mRNA entry channel. Unstructured mRNAs accommodate independently of eIF4E, whereas structured mRNAs require an upstream eIF4E-dependent step, enabling selective recruitment under limiting eIF4E. Arrested complexes can convert directly into the accommodated state without dissociation, revealing a reversible standby intermediate poised for activation. Together, our findings establish mRNA accommodation as a ribosome-intrinsic checkpoint governing initiation and provide a framework for selective translation.

## Introduction

Translation initiation begins with recruitment of mRNA to the small ribosomal subunit, a key step that determines which transcripts are selected for protein synthesis. Numerous studies have defined the roles of initiation factors, including the cap-binding complex eIF4F, in promoting mRNA recruitment^1,2^. It remains unclear how different mRNAs are selectively engaged at this stage. Current models emphasize differences in mRNA structure and competition for limiting initiation factors, yet these frameworks do not explain how the ribosome determines whether an engaged mRNA proceeds to productive accommodation and initiation or remains in a stable ribosome-bound intermediate state. In particular, whether mRNA recruitment proceeds through distinct ribosome-bound intermediates that govern commitment to productive accommodation has not been established, in part because previous studies have largely relied on stable intermediates that may obscure transient steps during mRNA engagement^3,4^.

Recent structural studies revealed an extended ∼70-nt mRNA path across the human 48S complex, far longer than the canonical ∼25-nt binding channel (Fig. S1). This architecture suggests that mRNA engagement with the ribosome likely involves multiple conformational rearrangements rather than a single binding event^5,6^. This extended path is largely formed by eIF3, with core domains of eIF4F positioned outside the exit channel, suggesting that mRNA is “slotted” rather than threaded into the entry channel^5^. Among these factors, the cap-binding complex eIF4F, comprising eIF4E, eIF4A, and eIF4G, plays a central role in mRNA recruitment. While eIF4A provides an ATP-dependent helicase activity together with eIF4G, eIF4E binds the 5′ cap and activates mRNA binding and helicase functions of eIF4F through interaction with eIF4G^7,8^. This interaction is tightly regulated through sequestration of eIF4E by 4E-BPs^9^. In addition, structural and biochemical studies suggest that the C-terminal tail (CTT) of eIF3j obstructs part of the mRNA path near the entry channel (Fig. S1), requiring displacement for full mRNA engagement^10–13^. Given that only single-stranded mRNA can be accommodated within the entry channel, these structural features further suggest that mRNA secondary structure must be resolved during engagement with the ribosome. However, how eIF3j kinetically regulates mRNA recruitment, and how these structural features contribute to selective mRNA engagement, remain unclear.

Here, we use real-time fluorescence assays and a fully reconstituted human initiation system to define the early steps of mRNA loading onto the 40S and 43S complexes. We find that mRNA recruitment proceeds through a branched kinetic pathway involving three conformational states intrinsic to the 40S: a rapid, dynamic sampling state; a productive accommodated state; and a long-lived arrested state. These states are shaped by mRNA structure, 40S conformation, eIF3, and eIF3j, which together regulate partitioning between productive accommodation and arrested state in the 43S complex. In contrast, eIF4F promotes accommodation in an ATP-dependent manner that is kinetically coupled to displacement of eIF3j from the entry channel. Accommodation of unstructured mRNA bypasses the need for eIF4E, whereas structured mRNAs require an upstream eIF4E-dependent activation step, revealing how the accommodation checkpoint can selectively partition mRNAs into distinct recruitment pathways under conditions of limiting eIF4E availability. We further show that mRNA in the arrested state is not a dead-end intermediate but can be directly converted into the accommodated state by eIF4F after binding to the 43S, suggesting that mRNA activation can occur after ribosome engagement and that the arrested state provides a kinetic window for regulating this transition. Together, these findings identify mRNA accommodation as a ribosome-intrinsic kinetic checkpoint that determines whether recruited mRNAs proceed toward productive initiation. This framework provides a mechanistic basis for selective mRNA recruitment and establishes early mRNA loading as a regulated, multistep conformational process rather than a simple binding event.

## Results

### mRNA undergoes a conformation change to two distinct states after rapid binding to the 40S or 43S

To track the early steps of mRNA engagement with the human 40S or 43S in real time, we developed a fluorescence anisotropy (FA) assay using 71-nt unstructured CAA repeat RNA fluorescently labeled at the 3′-end (CAA71-Fl, with no AUG codon), which shows significant increase in its FA (>0.1) upon binding to the ribosomes (Fig. 1A). This assay uses a reconstituted human initiation system capable of forming the 48S on start codons in a manner highly dependent on eIF4F (Figs. S2A-B), as consistent with previous studies using the same system^5,14^. Upon rapid mixing with the 43S in a stoppedlllflow fluorimeter, FA traces of CAA71-Fl consistently resolve into three distinct kinetic phases, indicating that the mRNA occupies three binding states (tentatively designated states 1, 2, and 3) differing by ≥5lllfold in their apparent rate constants (Fig. 1A). Interestingly, an addition of eIF4F increases the amplitude of the second phase while decreasing the third in a manner strongly dependent on ATP, suggesting that eIF4F promotes mRNA recruitment specifically as the state 2 (designated accommodated state).

**Figure 1.**
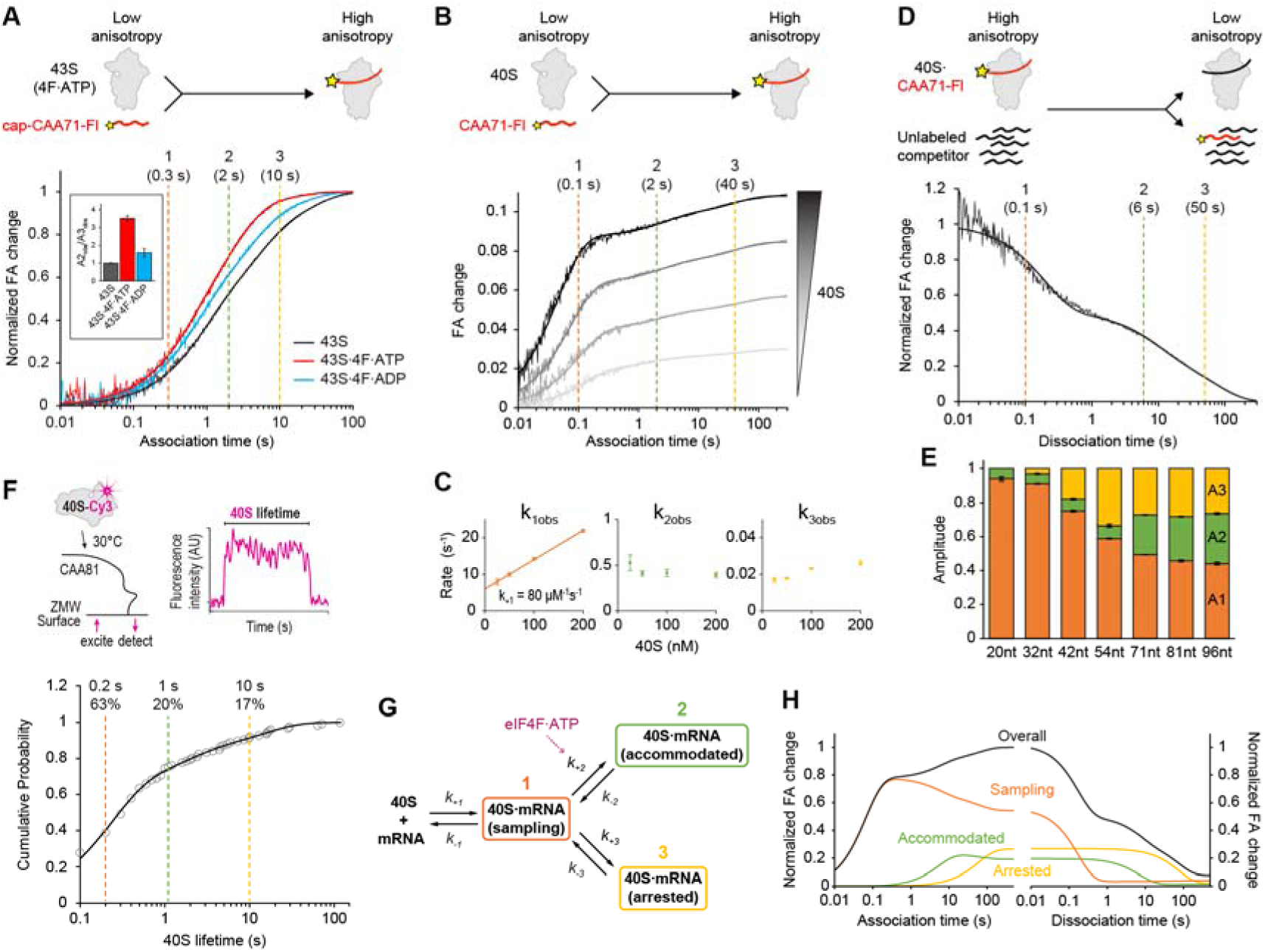
mRNA undergoes a conformation change to two distinct states after rapid binding to the ribosome. **(A)** Real-time association kinetics of cap-CAA71-Fl for the 43S complex in the presence or absence of eIF4F, monitored by fluorescence anisotropy (FA) change. Traces were fit to triple-exponential functions (thick lines), with half-times for each kinetic phase are indicated. Inset shows normalized ratios of the second- and third-phase amplitudes (A2_obs_/A3_obs_). **(B)** Association kinetics of CAA71-Fl with increasing concentrations of the 40S subunit in the absence of initiation factors, fit to triple-exponential functions (thick lines). **(C)** Apparent rate plots for each association phase in (B). **(D)** Dissociation kinetics of CAA71-Fl for the 40S subunit, fit to a triple-exponential function. **(E)** Amplitudes of dissociation phases for 3′-labeled CAA repeat RNAs of varying lengths (20–96 nt). **(F)** Schematic of the single-molecule real-time lifetime assay at 30°C, and a cumulative probability plot for 40S dissociation from CAA81, fit to a triple-exponential function (solid line) with half-lives and amplitudes indicated for each subpopulation. **(G)** Branched kinetic model of mRNA loading, showing rapid sampling (phase 1), followed by mRNA transition either into a productive accommodated state (phase 2) or a stable arrested state (phase 3). eIF4F promotes mRNA accommodation in an ATP-dependent manner. **(H)** Deconvolution of state subpopulations obtained from global fitting to association and dissociation dataset, using the branched model. All data are represented as mean ± SEM (n=3).

Association with the 40S in the absence of initiation factors also produces similar threelllphase kinetics, indicating that these states fundamentally originate from the 40S rather than initiation factors (Fig. 1B). A titration of the 40S suggests that state 1 is the concentration-dependent rapid binding step (designated sampling), while accommodated state and state 3 (designated arrested state) likely represent slower conformational transitions following initial binding (Fig. 1C).

Dissociation kinetics also exhibits the consistent tri-phasic behavior, confirming that the three association phases reflect genuine intermediates with distinct stabilities (Fig. 1D). We next examined how mRNA length affects the distribution among these states (Fig. S2D and Table 1). Despite similar apparent affinities for the 40S (Fig. S2C), RNAs 20–32 nt in length predominantly occupies sampling state (>90%, Fig. 1E), whereas those 42–54 nt increasingly populate arrested state (up to 34%). RNAs ≥71 nt show the highest level of accommodated state (23-30%) with slight reduction in arrested state (down to ∼27%), suggesting that a minimum of ∼70 nt is required to observe the full landscape of conformational transitions involved in early mRNA engagement in the absence of initiation factors. All states with CAA71-Fl could be equally competed by the same (71 nt) or short (20 nt, populating 94% in sampling state) unlabeled RNA (Fig. S2E), indicating that they share an overlapping 40S binding surface, consistent with accommodated and arrested states representing conformational rearrangements rather than distinct binding sites. A complementary single-molecule lifetime analysis of the 40S subunit on 81-nt CAA repeat RNA tethered on a surface corroborated the presence of these three states (Fig. 1F), although the rates are slightly faster likely due to a higher temperature (30°C instead of 25°C). The agreement between these independent measurements supports the conclusion that the observed multistate behavior reflects intrinsic features of mRNA recruitment rather than an artifact of bulk averaging.

Together, these results support a branched pathway in which mRNA, following initial sampling, partitions into either a productive accommodated state promoted by eIF4F or a long-lived arrested state, with all states being intrinsic to the 40S subunit (Fig. 1G). A model-based global fitting yields reasonable fit across the set of association and dissociation curves for 40S binding (Fig. S1G), enabling deconvolution of the conformation states (Fig. 1G). The rates obtained upon manual calculations (see methods) are well consistent with those determined in the global fitting (Table 2), and thus used in the following analysis for simplicity.

### mRNA structure and 40S conformation shape the conformational fate of mRNA

We next investigated regulatory features that bias mRNA toward either the accommodated or arrested pathway. Strikingly, a structured 76-nt β-globin 5′UTR RNA (hereafter βG-RNA) shows 4-fold slower sampling rate (consistent with 7-fold lower affinity for the 40S, Figs. 2A and S3A-C, Table 2), and is strongly biased toward the arrested state (69% versus 27% with CAA71) with no detectable amount of accommodated state in dissociation analysis (Figs. 2B-C), indicating that intrinsic mRNA structure itself can enforce arrested state. Alternatively, high magnesium (10 mM instead of 2 mM), which is known to induce a closed, inactive conformation of the 40S^15^, also makes CAA71 biased toward arrested state (64%, due to 4-fold faster forward and 3-fold slower backward rates, Figs. 2C-D and S3D). Meanwhile, it stabilizes sampling state by 5-fold and thereby increases affinity by ∼5-fold, consistent of narrowed mRNA binding cleft in closed 40S^16,17^ (Figs. 2A and S3A-B). Together, these results indicate that mRNA structure and 40S conformation strongly influence partitioning between accommodated and arrested states, favoring arrest under inhibitory conditions.

**Figure 2.**
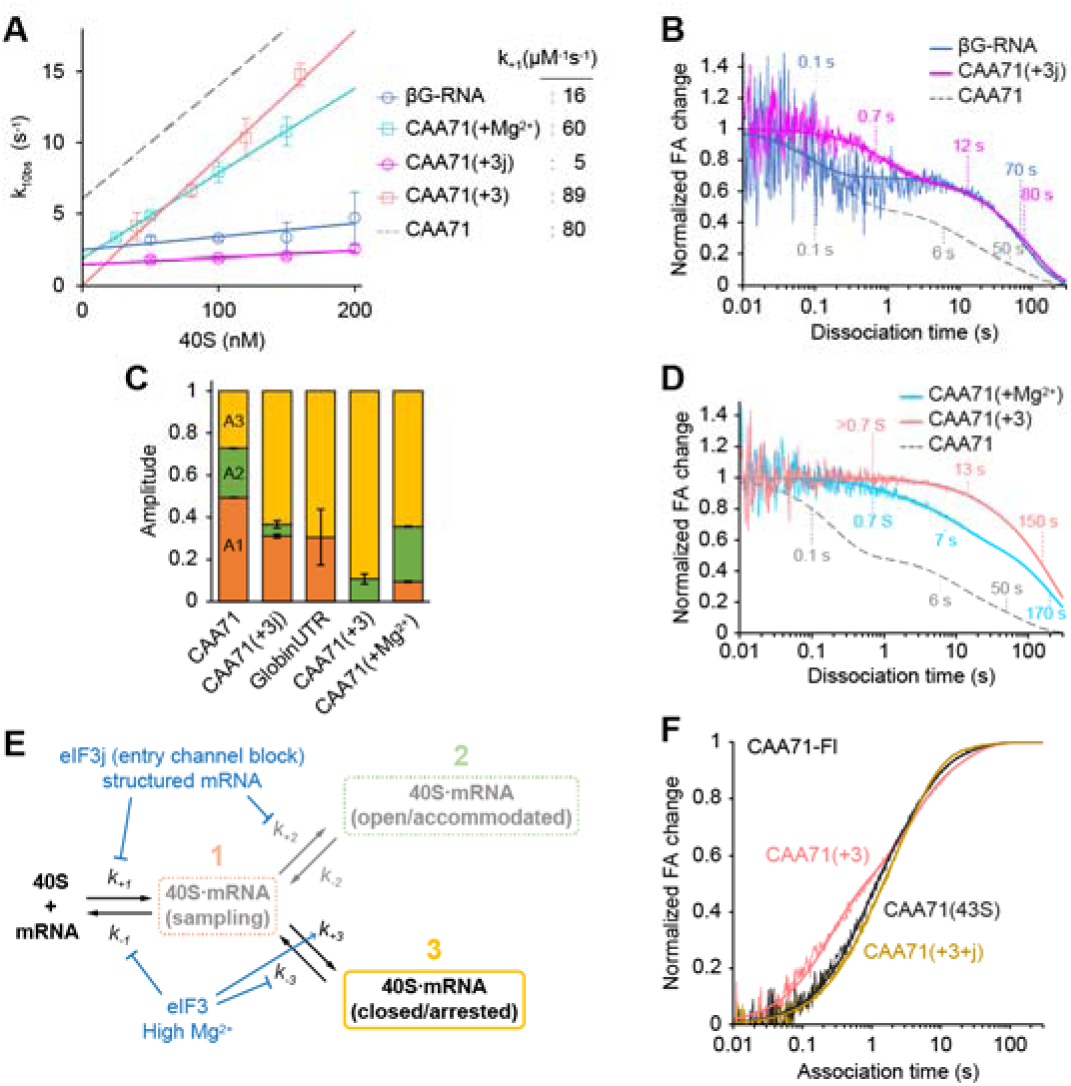
mRNA structure, 40S conformation, eIF3, and eIF3j shape the conformational fate of mRNA. **(A)** Apparent sampling rate plots for CAA71-Fl or βG-RNA-Fl measured under various conditions, including 40S with high magnesium (CAA71+Mg^2+^), 40S·eIF3j (CAA71+3j), and 40S·eIF3 (CAA71+3). **(B)** 40S dissociation kinetics for βG-RNA-Fl and for CAA71+3j. **(C)** Amplitudes of dissociation phases under the various conditions. **(D)** Dissociation kinetics for CAA71+Mg^2+^ and for CAA71+3. **(E)** Schematic summarizing effects of mRNA structure, high magnesium (closed 40S), eIF3, and eIF3j on mRNA kinetics, converging into the arrested state. **(F)** Association kinetics of CAA71-Fl for the 43S complex compared to 40S·eIF3 and 40S·eIF3·eIF3j. All data are represented as mean ± SEM (n=3).

### eIF3 and eIF3j promotes mRNA arrest in the 43S

Because eIF3j C-terminal tail (CTT) interferes with mRNA just outside the entry channel^13^ (Fig. S1), we tested whether it modulates mRNA kinetics. eIF3j strongly inhibits both sampling (by 16lllfold slower on rate but with 5-fold slower off rate, consistent with 5-fold reduction in affinity, Figs. 2A-B, and S3A), and accommodation (primarily by 5-fold slower forward rate), which leads to accumulation of CAA71 in the arrested state (63%, Fig. 2C). These effects closely resemble those with βG-RNA, and thus suggest that accommodation step involves engagement with mRNA entry channel, which requires displacement of the eIF3j-CTT and unstructured region in mRNA. In contrast to eIF3j, the remainder of eIF3 core complex increases affinity for the 40S by >5-fold (Fig. S3A), primarily through stabilizing the sampling state (Figs. 2A and S3B, see methods). Yet it also strongly favored the arrested state (89%, due to >5-fold faster forward and 3-fold slower backward rates, Figs. 2C-D) with minimal accommodated state (estimated to be ≤11%). These behaviors parallel the effects of high magnesium, suggesting that eIF3 also promotes closed 40S conformation, thereby facilitating mRNA arrest, besides extending mRNA path outside the entry and exit channels (Fig. S1). Together, eIF3 and eIF3j converge mechanistically to slow initial mRNA sampling, oppose productive accommodation, and stabilize mRNA in the arrested state (Fig. 2E). This trend persists up to the 43S (Fig. 2F), indicating that mRNA loading to the 43S is kinetically shaped by eIF3 and eIF3j toward the arrested state.

### mRNA structure partitions eIF4E-dependent and -independent pathways for accommodation

Having these kinetic models, we examined eIF4F-dependent steps in mRNA loading to the 43S. As the 43S is capable of downstream steps after mRNA loading (*e.g.* scanning), we only tested association kinetics with capped mRNAs harboring no AUG codon for physiological concentrations of 43S and eIF4F (60 nM each)^18^. For unstructured CAA71, eIF4F robustly favors accommodation without significantly changing the apparent rates (<2-fold) in a manner highly dependent on ATP but not eIF4E or 5’ cap (Figs. 3A-B and Table 3), suggesting that eIF4G and eIF4A together modulate ribosomal conformation, likely toward the dynamic open state, to facilitate accommodation rather than directly acting on mRNA binding and engagement. The structured βG-RNA shows ∼5-fold slower association in the absence of eIF4F, while eIF4F recovers sampling and accommodation rates by ∼4-fold, promoting accommodation over arrest, again in an ATP-dependent manner (Figs. 3C-D). In contrast to unstructured RNA, productive accommodation of structured RNA exhibits a strong dependence on eIF4E despite showing little dependence on the 5′ cap. These observations are consistent with an upstream eIF4E-dependent mRNA remodeling or restructuring step^7^ that precedes progression to the accommodated state. These findings reveal that eIF4F can overcome barriers imposed by mRNA structure, ribosome conformation, and associated initiation factors during accommodation. Structured mRNAs require an upstream eIF4E-dependent step, whereas unstructured mRNAs can bypass this requirement and proceed through an eIF4E-independent accommodation pathway (Fig. 3E).

**Figure 3.**
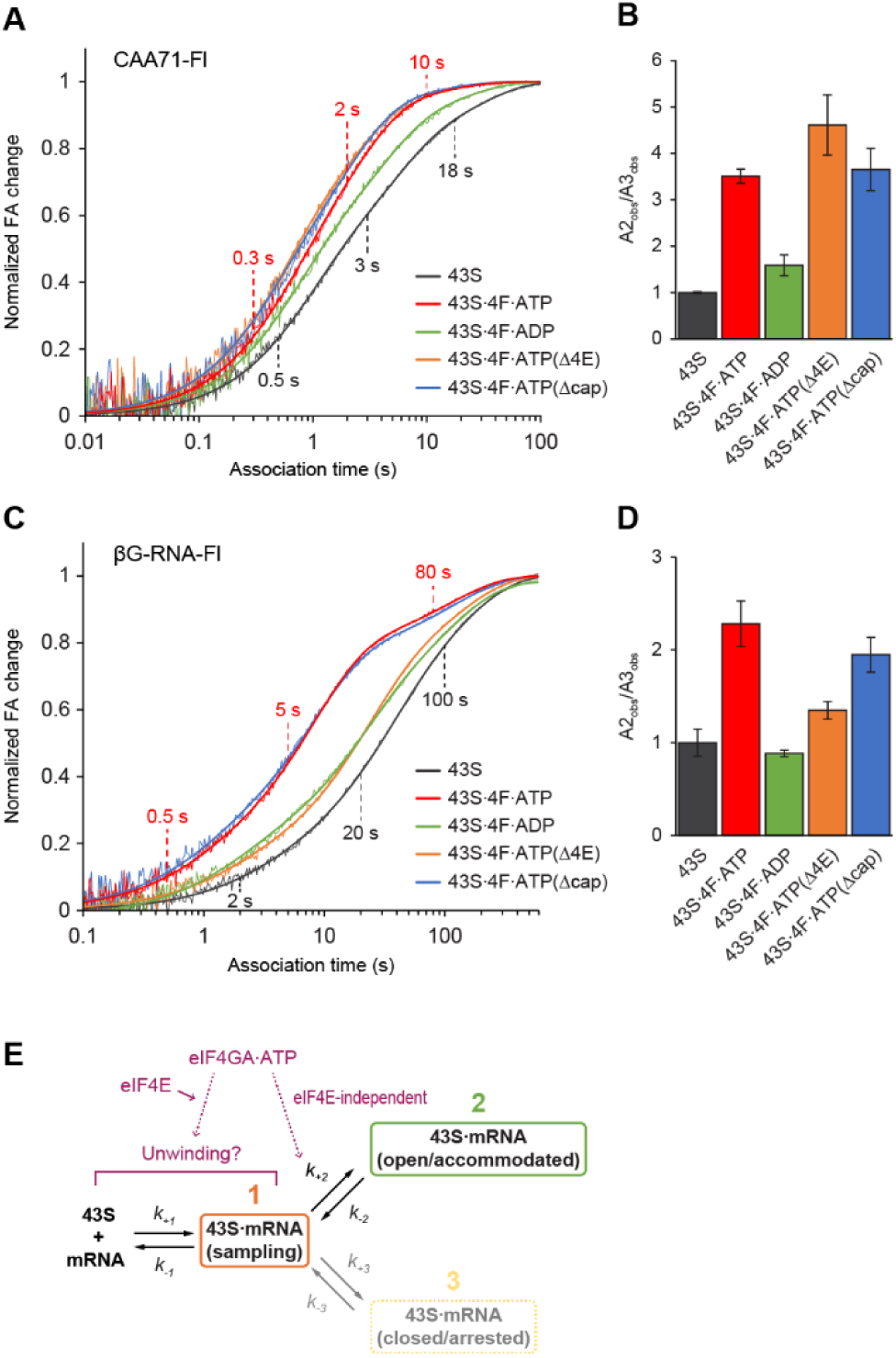
mRNA structure partitions eIF4E-dependent and -independent pathways for accommodation. **(A, C)** Association kinetics of CAA71-Fl or βG-RNA-Fl for the 43S complex in the presence or absence of eIF4F subunits. Half-times for each kinetic phase are indicated. **(B, D)** Normalized ratios of the second- and third-phase amplitudes (A2_obs_/A3_obs_). **(E)** Schematic summarizing the eIF4E-dependent and -independent effects of eIF4F on mRNA loading to the 43S. All data are represented as mean ± SEM (n=3).

### Displacement of eIF3j is kinetically coupled with mRNA accommodation

Because eIF4F and mRNA together cause a conformation change to destabilize eIF3j at the entry channel (from 3j_IN_ to 3j_OUT_ state, Figs. S1 and S4), thereby weakening affinity for the 43S by 10-fold^12^, we used FA signals from fluorescently labeled eIF3j (3j-Fl) to monitor this conformational change associated with accommodation. Under a limiting amount of the 43S (30 nM compared to *K*_d_ ≈ 140 nM as 3j_OUT_^12^), eIF3j dissociates from the 43S in a biphasic manner after rapid mixing with CAA71 in the presence of eIF4F. The release depends strongly on ATP with the fast phase (73%) being consistent with the rate of accommodation (0.61 s^-1^ versus 0.43 s^-1^, Fig. 4A and Table 4). Similar to mRNA kinetics, the fast and slow rates are slower when βG-RNA is used instead, while only the fast rate and amplitude are recovered by eIF4F in a both ATP- and eIF4E-dependent manner (Fig. 4B). These results strongly support a tight mechanistic link between mRNA accommodation and eIF3j release, and further indicate that progression to accommodation for structured mRNAs is limited by the upstream eIF4E-dependent step. The slow-phase rates closely match the arrest rates for both structured and unstructured mRNAs. Notably, the slow phase amplitude for structured mRNA increases in an ATP-dependent manner (Fig. 4B), suggesting that mRNA structure increases a subpopulation of mRNA that can initially be arrested and then be activated into the accommodated state by eIF4F and ATP. The structured mRNA preferentially follows this pathway especially when eIF4E is depleted (73% subpopulation, Fig. 4B).

### Arrested complexes can be directly activated by eIF4F into the accommodated state

To directly test whether the arrested complexes can be converted into the accommodated state, CAA71 was pre-loaded onto the 43S (+3j-Fl) in the presence of eIF4F but absence of ATP, conditions that favor accumulation of the arrested state. ATP was then added to trigger accommodation. Following a rapid minor increase in 3j-Fl anisotropy (0.79 s^-1^) likely reflecting a conformational rearrangement preceding eIF3j release, the majority of eIF3j release (82%) occurred with a rate more than 10-fold faster than the intrinsic backward rate of the arrested state (0.28 s⁻¹ versus <0.03 s⁻¹; Fig. 4C). Thus, the observed transition cannot be explained by mRNA first escaping the arrested state and returning to the sampling state. Instead, these results demonstrate that arrested complexes can be directly and rapidly converted into the accommodated state by eIF4F and ATP while mRNA remaining associated with the ribosome (Fig. 4D). Together, these findings establish that the arrested state is not a kinetic dead end but a reversible ribosome-bound intermediate poised for productive accommodation.

**Figure 4.**
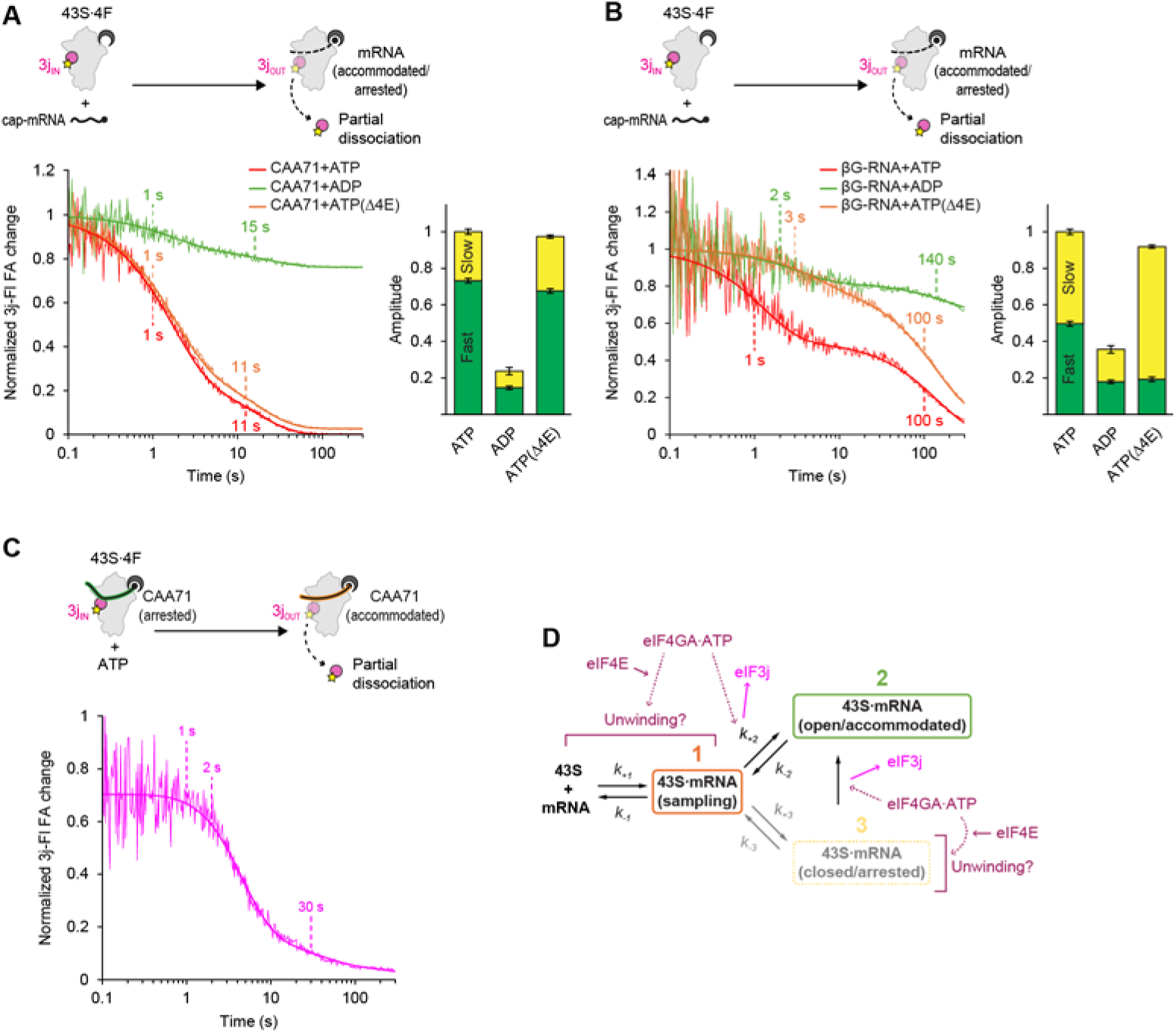
Displacement of eIF3j is kinetically coupled with mRNA accommodation. **(A, B)** Release of 3j-Fl from the 43S·eIF4F complex triggered by cap-CAA71 or cap-βG-RNA, monitored by FA change. Curves are fit to double-exponential functions (thick lines), with half-times for each phase indicated. Bar graphs show amplitudes of the fast and slow phases. **(C)** Release of 3j-Fl from 43S·eIF4F preloaded with cap-CAA71 in the arrested state (assembled without ATP), triggered by ATP. A curve is fit to a triple-exponential function (thick lines), with half-times for each phase indicated. **(D)** Schematic summarizing the mechanism of eIF3j release during two possible pathways of mRNA accommodation. eIF4F and mRNA cooperatively drive a conformational transition of the 43S that displaces the eIF3j-CTT from the entry channel. This displacement is tightly coupled to eIF4F- and ATP-dependent mRNA accommodation, which may be preceded by eIF4F-dependent unwinding. All data are represented as mean ± SEM (n=3).

## Discussion

Here, we define mRNA recruitment as a multistep, branched process intrinsic to the ribosome, in which mRNAs partition between arrested and accommodated states following an initial sampling step. This identifies mRNA accommodation as a key kinetic checkpoint that determines whether ribosome engagement proceeds to initiation. In this framework, mRNA structure and initiation factors do not simply influence binding, but instead regulate progression through ribosome-bound intermediates, thereby shaping the outcome of mRNA recruitment.

The sampling step likely represents a transient interaction in which mRNA rapidly associates with and dissociates from the ribosome. Short unstructured RNAs remain almost exclusively in this state, consistent with rapid loading into the canonical ∼25-nt binding channel as observed in yeast reconstitution systems^19^. In contrast, longer RNAs (>70 nt) populate both the accommodated and arrested states, indicating that a minimum RNA length is required to stabilize interactions extending beyond the mRNA channel and into adjacent ribosomal regions. The accommodated state is likely correlated with mRNA engagement into the entry channel and the open 40S conformation poised for downstream scanning. Its characteristic rate (≥0.1 s⁻¹) aligns well with the fastest estimates of initiation kinetics (∼4 s total)^20^, supporting the view that accommodation constitutes a key rate-limiting and regulatory step in the transition toward scanning. By contrast, the arrested state is associated with the closed 40S conformation with the narrowed mRNA channel, stabilizing mRNA but inhibiting scanning. While the inherent ability of the 40S to sample accommodated and arrested states suggests spontaneous fluctuations between open and closed conformations^15^, the arrested state is favored under conditions that restrict RNA movement within the mRNA path, such as structure within mRNA, elevated magnesium, or the presence of eIF3 and eIF3j.

Our data place eIF4F as a key regulator of progression through the ribosomal kinetic checkpoint that governs the transition from initial mRNA engagement to accommodation, acting to resolve barriers that otherwise bias mRNAs toward the arrested state. eIF4F resolves structural and ribosomal barriers to mRNA accommodation through both eIF4E-dependent and eIF4E-independent activities. The eIF4E-dependent activity is consistent with mRNA activation in canonical models, in which eIF4F binding to the 5′-end of mRNA promotes unwinding of nearby structure^1,2^. However, at the concentration of eIF4F used here (60 nM), eIF4F alone is unlikely to support substantial unwinding before 40S binding^7^. Thus, whether this eIF4E-dependent step occurs before ribosome binding, during engagement with the 43S, or both remains to be determined (Fig. 5). Complementary single-molecule studies suggest that eIF4F can first bind and remodel mRNA to promote subsequent 43S loading, particularly under conditions where eIF4F is readily available while the 43S is limited ^21^. Our results are consistent with a pathway in which mRNAs can engage the ribosome before productive eIF4F-dependent activation, entering ribosome-bound intermediates whose progression is governed by the accommodation checkpoint. These observations suggest that eIF4F-first and ribosome-first pathways may represent alternative routes that converge on a common accommodation checkpoint, with their relative use determined by mRNA structure and the availabilities of eIF4F (or eIF4E) and the 43S. For example, the presence of 5′-cap stabilizes eIF4F on mRNA and promotes more efficient unwinding in the presence of eIF4B and ATP in yeast^22^, highlighting conditions under which pre-binding activation is favored. The eIF4E-independent activity of eIF4F does not appreciably accelerate sampling of unstructured RNAs, which is consistent with the direct interaction between eIF4F and mRNA being highly dependent on eIF4E^8^, suggesting that it acts after the sampling step, likely through remodeling of the 43S. Consistent with this framework, mRNA can remain ribosome-bound in the absence of eIF4F activity, as observed in yeast where many transcripts remain associated with ribosomes upon depletion of eIF4G despite strong inhibition of translation^23^. This framework suggests that mRNAs with less structured 5′ UTRs may be preferentially recruited under conditions of limiting eIF4E availability, consistent with selective translation programs such as heat shock^24^. This eIF4E-independent activity further suggests that eIF4F orthologs lacking eIF4E, such as eIF4G2^25,26^, may function primarily by promoting mRNA accommodation through regulation of the ribosomal checkpoint.

**Figure 5.**
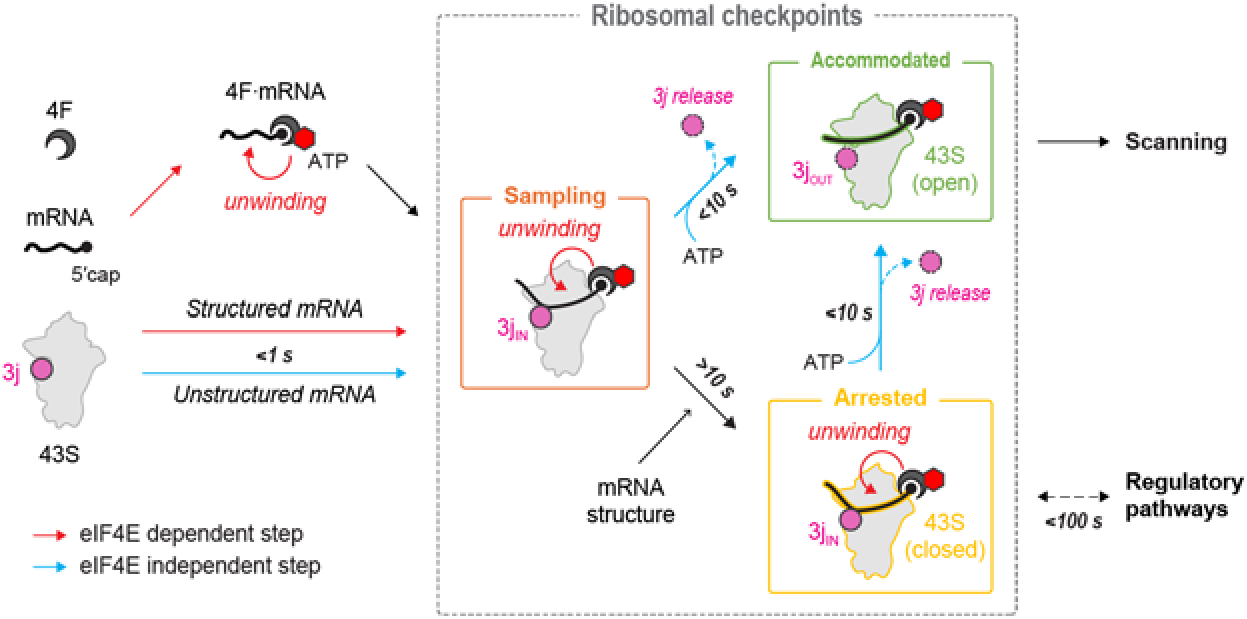
Proposed model for mRNA recruitment and its regulation. mRNA recruitment begins through two alternative pathways: direct binding of eIF4F to the mRNA 5′ end or independent association of mRNA and eIF4F with the 43S, depending on their relative availability. In both cases, mRNA rapidly binds the 43S and enters a transient sampling state. Unstructured mRNAs can bypass eIF4E at this stage, whereas structured mRNAs depend strongly on eIF4E likely to promote mRNA remodeling before and/or after ribosome binding, which may therefore be more sensitive to 4E-BPs and eIF4E phosphorylation^37,38^. After sampling, mRNAs either proceed into a productive accommodated state or transition into a slowly formed, long-lived arrested state. Accommodation occurs over several seconds and is driven by ATP-dependent but eIF4E-independent activity of eIF4F, accompanied by 40S opening and displacement of eIF3j-CTT from the entry channel. Arrested complexes (∼100 s lifetime) are reversible intermediates that can be activated by eIF4F once inhibitory barriers are resolved, for example by eIF4F itself or by accessory helicases including DDX3^39–42^ and DHX29^43,44^. Structured 5′ UTRs, depletion of eIF4F, and entry channel binding factors (such as eIF3j, Nsp1, and PDCD4) favor arrest, which may lead to downstream regulatory pathways such as stress granule formation.

Our data show that the arrested state is not a kinetic dead end but a reversible intermediate that can be directly converted into the accommodated state on the ribosome. The rapid conversion of the arrested state to the accommodated state upon ATP addition, coupled with displacement of eIF3j from the entry channel, indicates that this transition can occur directly on the ribosome without complete mRNA dissociation. This suggests that the arrested state functions as a standby-like intermediate, allowing mRNAs to remain associated with the ribosome while awaiting productive accommodation and subsequent scanning. Our data establish a kinetic window—on the order of tens of seconds under our experimental conditions—during which arrested complexes remain competent for subsequent accommodation, possibly providing an opportunity for barriers to accommodation to be overcome (Fig. 5). This framework may help explain how stalled pre-initiation complexes, such as those in stress granules, remain poised for rapid reactivation when initiation factors become available^27^.

As eIF3j displacement depends on eIF4F, eIF2, and the presence of a single-stranded region in mRNA^12^, our study highlights a role of eIF3j as a gatekeeper of the entry channel that limits nonspecific mRNA accommodation. In this model, eIF3j prevents premature engagement of mRNA with the ribosome, thereby enforcing productive eIF4F-dependent accommodation and subsequent recruitment to the start codon. This mechanism may be particularly important for regulating the recruitment of unstructured mRNAs, which might bypass eIF4F-dependent control^4^ (Fig. S2A). We further propose that this mechanism may represent a broader class of entry channel regulatory factors, as several translation regulators, including coronavirus Nsp1^28,29^, PDCD4^30,31^, SERBP1^32^, and LARP1^33,34^, have been reported to interfere with the mRNA entry channel (Fig. 5).

Together, our findings reveal that mRNA structure, 40S conformation, eIF3j, eIF3, and eIF4F collectively define a kinetic checkpoint that governs progression to productive accommodation and subsequent scanning. Unstructured mRNAs can undergo eIF4E-independent accommodation, potentially enabling their preferential translation when eIF4E is limited, whereas structured mRNAs require eIF4E-dependent remodeling to achieve accommodation. Our data further establish that mRNA recruitment proceeds through a reversible ribosome-bound intermediate that can be directly converted into the accommodated state, identifying mRNA accommodation as a ribosome-intrinsic kinetic checkpoint that determines whether recruited mRNAs proceed toward productive initiation. More broadly, this framework provides a quantitative and mechanistic basis for how ribosomes discriminate among mRNAs before scanning, reframing early translation initiation as a regulated, multistep conformational decision rather than a simple binding event (Fig. 5).

## Methods

### Sample preparations

Human initiation factors and the 40S subunit were prepared as previously described^7,10,14,35^. eIFs 1, 1A, 4E, 4A1, and 5 were purified from overexpressing *Escherichia coli* BL21 (DE3), while eIF3j (and its variant), eIF4B, and eIF4G1 (residues 165-1599) were purified from overexpressing Sf9 insect cells. Native eIF2, eIF3 (lacking eIF3j), and the 40S subunit were purified from HeLa cells. The eIF2-TC was reconstituted with GMPPNP and in vitro transcribed tRNA_i_ methionylated with *E. coli* methionyl-tRNA synthetase. Fluorescently labeled eIF3j (3j-Fl) was generated by labeling T235C single cysteine mutant with fluorescein-5-maleimide as previously described^12^. mRNAs were transcribed using T7 RNA polymerase with synthetic oligo DNAs templates and labeled as previously described^10,12^. The sequences of mRNAs are summarized in Table 5. Model β-globin 5′UTR mRNA was refolded at 1 μM in buffer A (20 mM tris-HCl pH 7.5, 70 mM KCl, 2 mM MgCl_2_, 0.1 mM spermidine, 2 mM DTT, and 5% glycerol) by incubation at 70°C for 2 min, followed by slow cooling (−1°C/min) to 25°C in a thermal cycler. The RNA was then concentrated using Amicon Ultra device (3kDa cutoff, Millipore). The 3′ ends of mRNAs were labeled with fluorescein-5-thiosemicarbazide after oxidization. The 5′ end of CAA71 was labeled with fluorescein-5-maleimide, after dephosphorylation with calf intestinal phosphatase and re-phosphorylation with ATPγS using T4 polynucleotide kinase. Both labeled and unlabeled mRNAs were capped using a faustovirus capping enzyme (New England Biolabs), and purified through a Micro Bio-Spin P-6 column.

### Ribosomal complex formation

2×43S complexes were assembled by mixing 120 nM 40S, 150 nM eIF3, 150 nM eIF2-TC, 250 nM each of eIFs 1, 1A, 3j, and 5, with or without 120 nM eIF4G1, 250 nM each of eIFs 4A1, 4E, and 4B in buffer B (20 mM tris-HCl pH 7.5, 40 mM KCl, 60 mM potassium acetate, 2.2 mM magnesium acetate, 0.1 mM spermidine, 2 mM DTT, and 5% glycerol) supplemented with 0.5 mM ATP·Mg or ADP·Mg, followed by incubation at 37°C for 10 min. For the eIF3j release assays, the complexes were prepared analogously but using 60 nM 40S and 40 nM 3j-Fl instead of unlabeled eIF3j.

### Start site recognition assays

Reactions were initiated by mixing 2×43S with 30 nM cap-CAA81(AUG)-Fl in buffer B at 30°C. For RelE cleavage assays, 10 μl reaction mixture was incubated at 37°C for 10 min with 2 μl mixture of 48 μM RelE and 60 μM unlabeled cap-CAA81(AUG), quenched with 12 μl of 10M Urea, and analyzed by 10% urea-PAGE as previously described^5^. For the FA assays, start codon recognition was monitored via enhanced binding stability of mRNA on the 43S upon start codon recognition. Bound mRNA was competed with excess unlabeled mRNA competitor (13.3 μM) in a 384-well plate at 25°C for 1 hour. mRNA remain bound to the 43S was quantitated by FA change measured in a CLARIOstar Plus plate reader (BMG Labtech) using 482-16/530-40 filter set with polarizers. The assay was validated to be specific to AUG codon (Fig. S2A), and compared with RelE cleavages (Fig. S2B).

### Measurements of mRNA binding kinetics

For the association kinetics, varying concentrations of 40S were preincubated with or without 350 nM eIF3 and/or 2 μM eIF3j in buffer A at 37°C for 10 min. The high magnesium experiments used 10 mM MgCl_2_ instead of 2 mM. The preincubated ribosomes were loaded to AutoSF-120 stopped-flow fluorimeter (Kintek) at 25°C, further incubated for >5 min, and rapidly mixed with 40 nM 3′-labeled mRNA (20 μl + 20 μl per shot). A time-dependent changes in FA were monitored with log-scale intervals using 480 nm emission light (20 nm monochromator slits) and 531/40 excitation filter with polarizers. Dissociation kinetics were measured analogously, except labeled mRNA was prebound to ribosomal complexes, then competed with 6 μM unlabeled competitor RNA. Three sets of independent experiments (5-13 shots each) were performed for all conditions.

### Global fitting

A set of association curves and dissociation curve for the 40S were globally fit using Kintek Explorer 6.3^36^, based on the branched model (Fig. S2G and Tabel 2). Consistent with no significant difference among dissociation curves for 3’-labeled and 5’labeled CAA71 (Fig. S2F), the global fitting confirms that FA changes caused by conformation changes are minimal (<10% compared to the FA change upon binding), suggesting that FA amplitudes directly reflect the population of corresponding states. The resulting deconvolution of sampling, accommodated, and arrested states align with the triple-exponential fit to the dissociation curve, whereas later phases of association curves reflect mixtures of states that lead to small signals, making them less reliable for rate extraction (Fig. 1H).

### Kinetic analysis

FA traces were fitted with sigma-weighted double- or triple-exponential functions using Kintek software or KareidaGraph 4.0. In dissociation analysis, the amplitudes (A1, A2, and A3) and rate constants (k_-1_, k_-2_, and k_-3_) were obtained directly from fitting. Sampling rates (k_+1_) were obtained from the observed association rate (k_1obs_) plots, which also provided estimates for k_-1_ (termed k_-1_*) well consistent with the k_-1_ obtained from dissociation analysis, indicating that competitor RNA did not cause facilitated dissociation in the dissociation experiments (Table 2). Accommodation (k_+2_) and arrest (k_+3_) rate constants were calculated from amplitude ratios using k_+2_/k_-2_ = A2/A1 or k_+3_/k_-3_ = A3/A1, according to the branched model. For CAA71 binding to 40S·eIF3 (CAA71(+3)), dissociation kinetics follows a double exponential function, while association remain to be triphasic with no significant shifts in apparent rates, suggesting that off rates of sampling and accommodated states become indistinguishable. Accordingly, k_-1_* was used as an estimate for k_-1_, while an apparent amplitude of the fast phase (11%) was considered to be sum of A1 and A2 (see Table 2).

### Single molecule analysis

3′-biotinylated CAA81 mRNA and Cy3-labeled human 40S ribosomal subunits (from HEK 293T cells expressing ybbR-tagged RPS15) were prepared as previously described^14^. The real-time single-molecule assay was conducted at 30 °C on a modified Pacific Biosciences RSII DNA sequencer using Maggie software (v. 2.3.0.3.154799) as previously described^14^. Briefly, biotinylated mRNA was tethered to a neutravidin-coated zero-mode waveguide (ZMW) chip surface. Data acquisition began by adding Cy3-40S subunits in imaging buffer (20LmM HEPES–KOH pH 7.5, 70 mM potassium acetate, 2.5 mM magnesium acetate, 0.25LmM spermidine, 0.2 mg/ml creatine phosphokinase, 2 mM protocatechuic acid, and 0.06 U/µl protocatechuate-3,4-dioxygenase) to the chip for a final concentration of 5 nM. The surface was excited with a 532 nm laser (0.32 µW µm⁻²), and Cy3 fluorescence emission was recorded at 10 frames per second for 900 seconds. Experimental movies captured on the PacBio RS II were processed on MATLAB 2024a as previously described^14^. Although many data points (28%) are accumulated at the shortest measurable lifetime due to the relatively slow record framerate (0.1 s), which could cause an underestimation of the fastest rate, the data was well-fit to a triple exponential cumulative distribution function to obtain dissociation rate constants.

### eIF3j kinetics

eIF3j release was monitored similarly to mRNA loading kinetics but using 3j-Fl. 2×43S (containing 3j-Fl) were rapidly mixed with 200 nM unlabeled cap-CAA71 or cap-βG-RNA. For pre-arrested complexes, 2×43S was preloaded with 200 nM cap-CAA71 without ATP, then mixed with 1 mM ATP·Mg to trigger accommodation. Dissociation of 3j-Fl from either the 43S or the 48S complex was measured by competition with 2 μM unlabeled eIF3j. Three sets of independent experiments (5-10 shots each) were performed for all conditions.

## Supporting information

Suppelementary Figures S1-4 and Tables 1-5

## Acknowledgements

We are grateful to members of the Fraser and Puglisi labs for helpful guidance, discussions, and feedback. We dedicate this paper to the memory of our long-time friend and colleague Nancy Villa, who passed away in February 2026. C.A. is supported by the Howard Hughes Medical Institute Gilliam Fellows Program and a Stanford Bio-X fellowship. This work was funded, in part, by the NIH (R35 GM152137 to C.S.F., R35 GM145306 and R01 AG064690 to J.D.P., K99 GM144678 to C.P.L.), the University of California Cancer Research Coordinating Committee grant (C26CR10046 to C.S.F.), and the Chan Zuckerberg Biohub Investigator Award (to J.D.P.).

## Author contributions

Conceptualization, M.S. and C.S.F.; Bulk kinetics and biochemistry: Methodology, M.S. and C.S.F.; Resources, M.S., N.V. and C.S.F.; Investigation, M.S.; Formal Analysis, M.S.; Validation, M.S.; Visualization, M.S.; Single molecule analysis: Methodology, C.A., C.P.L. and J.D.P.; Resources, C.A. and C.P.L.; Investigation, C.A. and C.P.L.; Formal analysis, C.A.; Validation, C.A.; Visualization, C.A.; Funding acquisition, C.P.L., J.D.P. and C.S.F.; Project administration, J.D.P. and C.S.F.; Supervision, J.D.P. and C.S.F.; Writing—original draft, M.S.; Writing—review and editing, M.S., C.A., C.P.L., J.D.P. and C.S.F.

## References

1. Sokabe, M. & Fraser, C. S. Toward a Kinetic Understanding of Eukaryotic Translation. Cold Spring Harb. Perspect. Biol. 11, (2019).

2. Brito Querido, J., Díaz-López, I. & Ramakrishnan, V. The molecular basis of translation initiation and its regulation in eukaryotes. Nature Reviews Molecular Cell Biology 2023 25:3 25, 168–186 (2023).

3. Andreev, D. E., Tierney, J. A. S. & Baranov, P. V. Translation Complex Profile Sequencing Allows Discrimination of Leaky Scanning and Reinitiation in Upstream Open Reading Frame-controlled Translation. J. Mol. Biol. 436, 168850 (2024).

4. Pestova, T. V & Kolupaeva, V. G. The roles of individual eukaryotic translation initiation factors in ribosomal scanning and initiation codon selection. Genes Dev. 16, 2906–22 (2002).

5. Brito Querido, J., et al. Structure of a human 48S translational initiation complex. Science 369, 1220–1227 (2020).

6. Brito Querido, J., et al. The structure of a human translation initiation complex reveals two independent roles for the helicase eIF4A. Nature Structural & Molecular Biology 2024 31:3 31, 455–464 (2024).

7. Feoktistova, K., Tuvshintogs, E., Do, A. & Fraser, C. S. Human eIF4E promotes mRNA restructuring by stimulating eIF4A helicase activity. Proc. Natl. Acad. Sci. U. S. A. 110, 13339–44 (2013).

8. Izidoro, M. S., Sokabe, M., Villa, N., Merrick, W. C. & Fraser, C. S. Human eukaryotic initiation factor 4E (eIF4E) and the nucleotide-bound state of eIF4A regulate eIF4F binding to RNA. Journal of Biological Chemistry 298, 102368 (2022).

9. Pause, A. et al. Insulin-dependent stimulation of protein synthesis by phosphorylation of a regulator of 5’-cap function. Nature 1994 371:6500 371, 762–767 (1994).

10. Fraser, C. S., Berry, K. E., Hershey, J. W. B. & Doudna, J. A. eIF3j is located in the decoding center of the human 40S ribosomal subunit. Mol. Cell 26, 811–9 (2007).

11. Fraser, C. S., Hershey, J. W. B. & Doudna, J. a. The pathway of hepatitis C virus mRNA recruitment to the human ribosome. Nat. Struct. Mol. Biol. 16, 397–404 (2009).

12. Sokabe, M. & Fraser, C. S. A helicase-independent activity of eIF4A in promoting mRNA recruitment to the human ribosome. Proc. Natl. Acad. Sci. U. S. A. 114, 6304–6309 (2017).

13. Kratzat, H. et al. A structural inventory of native ribosomal ABCE1-43S pre-initiation complexes. EMBO J. 40, EMBJ2020105179-(2021).

14. Lapointe, C. P. et al. eIF5B and eIF1A reorient initiator tRNA to allow ribosomal subunit joining. Nature 2022 607:7917 607, 185–190 (2022).

15. Shenvi, C. L., Dong, K. C., Friedman, E. M., Hanson, J. A. & Cate, J. H. D. Accessibility of 18S rRNA in human 40S subunits and 80S ribosomes at physiological magnesium ion concentrations—implications for the study of ribosome dynamics. RNA 11, 1898–1908 (2005).

16. Llácer, J. L. et al. Conformational Differences between Open and Closed States of the Eukaryotic Translation Initiation Complex. Mol. Cell 59, 399–412 (2015).

17. Yi, S. H. et al. Conformational rearrangements upon start codon recognition in human 48S translation initiation complex. Nucleic Acids Res. 50, 5282–5298 (2022).

18. Kumar, P., Hellen, C. U. T. & Pestova, T. V. Toward the mechanism of eIF4F-mediated ribosomal attachment to mammalian capped mRNAs. Genes Dev. 30, 1573–88 (2016).

19. Maag, D., Fekete, C. A., Gryczynski, Z. & Lorsch, J. R. A conformational change in the eukaryotic translation preinitiation complex and release of eIF1 signal recognition of the start codon. Mol. Cell 17, 265–75 (2005).

20. Shah, P., Ding, Y., Niemczyk, M., Kudla, G. & Plotkin, J. B. Rate-limiting steps in yeast protein translation. Cell 153, 1589–601 (2013).

21. Carlos Alvarado, C. P. L. J. W. M. S. R. G. A. A. D. C. I. S. M. Z. P. A. M. J. J. C. S. F. J. D. P. Remodeling of mRNA by eIF4F in human translation initiation. Manuscript submitted for publication (2026).

22. Gentry, R. C. et al. The mechanism of mRNA cap recognition. Nature 637, 736 (2024).

23. Park, E. H., Zhang, F., Warringer, J., Sunnerhagen, P. & Hinnebusch, A. G. Depletion of eIF4G from yeast cells narrows the range of translational efficiencies genome-wide. BMC Genomics 12, 68-(2011).

24. Hess, M. A. & Duncan, R. F. Sequence and structure determinants of Drosophila Hsp70 mRNA translation: 5’UTR secondary structure specifically inhibits heat shock protein mRNA translation. Nucleic Acids Res. 24, 2441–2449 (1996).

25. Imataka, H., Olsen, H. S. & Sonenberg, N. A new translational regulator with homology to eukaryotic translation initiation factor 4G. The EMBO Journal 1997 16:4 16, 817–825 (1997).

26. Yamanaka, S., Poksay, K. S., Arnold, K. S. & Innerarity, T. L. A novel translational repressor mRNA is edited extensively in livers containing tumors caused by the transgene expression of the apoB mRNA-editing enzyme. Genes Dev. 11, 321–333 (1997).

27. Buchan, J. R. & Parker, R. Eukaryotic Stress Granules: The Ins and Out of Translation. Mol. Cell 36, 932 (2009).

28. Schubert, K. et al. SARS-CoV-2 Nsp1 binds the ribosomal mRNA channel to inhibit translation. Nature Structural & Molecular Biology 2020 27:10 27, 959–966 (2020).

29. Thoms, M. et al. Structural basis for translational shutdown and immune evasion by the Nsp1 protein of SARS-CoV-2. Science (1979). 369, 1249–1256 (2020).

30. Ye, X. et al. Human tumor suppressor PDCD4 directly interacts with ribosomes to repress translation. Cell Res. 34, 522–525 (2024).

31. Brito Querido, J., et al. Human tumor suppressor protein Pdcd4 binds at the mRNA entry channel in the 40S small ribosomal subunit. Nat. Commun. 15, (2024).

32. Brown, A., Baird, M. R., Yip, M. C. J., Murray, J. & Shao, S. Structures of translationally inactive mammalian ribosomes. Elife 7, (2018).

33. Wolin, E. et al. SPIDR enables multiplexed mapping of RNA-protein interactions and uncovers a mechanism for selective translational suppression upon cell stress. Cell 188, 5384–5402.e25 (2025).

34. Saba, J. A. et al. LARP1 binds ribosomes and TOP mRNAs in repressed complexes. EMBO J. 43, 6555 (2024).

35. Sokabe, M. & Fraser, C. S. Human Eukaryotic Initiation Factor 2 (eIF2)-GTP-Met-tRNAi Ternary Complex and eIF3 Stabilize the 43S Preinitiation Complex. J. Biol. Chem. 289, 31827–31836 (2014).

36. Johnson, K. A., Simpson, Z. B. & Blom, T. Global kinetic explorer: a new computer program for dynamic simulation and fitting of kinetic data. Anal. Biochem. 387, 20–29 (2009).

37. Flynn, A. & Proud, C. G. Serine 209, not serine 53, is the major site of phosphorylation in initiation factor eIF-4E in serum-treated Chinese hamster ovary cells. J. Biol. Chem. 270, 21684–21688 (1995).

38. Bonneau, A. M. & Sonenberg, N. Involvement of the 24-kDa cap-binding protein in regulation of protein synthesis in mitosis. Journal of Biological Chemistry 262, 11134–11139 (1987).

39. Guenther, U. P. et al. The helicase Ded1p controls use of near-cognate translation initiation codons in 5′ UTRs. Nature 2018 559:7712 559, 130–134 (2018).

40. Zhou, F. et al. Transcriptome-wide analysis of the function of Ded1 in translation preinitiation complex assembly in a reconstituted in vitro system. Elife 13, (2024).

41. Soto-Rifo, R. et al. DEAD-box protein DDX3 associates with eIF4F to promote translation of selected mRNAs. EMBO J. 31, 3745 (2012).

42. Calviello, L. et al. DDX3 depletion represses translation of mRNAs with complex 5′ UTRs. Nucleic Acids Res. 49, 5336–5350 (2021).

43. Pisareva, V. P., Pisarev, A. V., Komar, A. A., Hellen, C. U. T. & Pestova, T. V. Translation Initiation on Mammalian mRNAs with Structured 5′UTRs Requires DExH-Box Protein DHX29. Cell 135, 1237–1250 (2008).

44. Hashem, Y. et al. Structure of the mammalian ribosomal 43S preinitiation complex bound to the scanning factor DHX29. Cell 153, 1108–19 (2013).

45. Meng, E. C. et al. UCSF ChimeraX: Tools for structure building and analysis. Protein Sci. 32, (2023).

46. Mathews, D. H. et al. Incorporating chemical modification constraints into a dynamic programming algorithm for prediction of RNA secondary structure. Proc. Natl. Acad. Sci. U. S. A. 101, 7287–7292 (2004).

