## Supplementary material for "A ribosomal kinetic checkpoint governs selective mRNA recruitment": Suppelementary Figures S1-4 and Tables 1-5

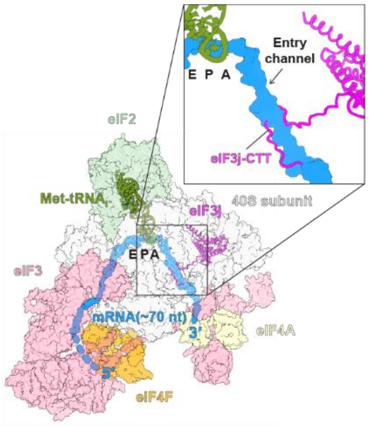


**Figure S1. Structure of the human 48S complex.**
The extended ~70 nt mRNA path is highlighted in blue (PDB 8OZ0[6]). The eIF3j (HCR1) C-terminal tail from the yeast 43S·ABCE1 structure (PDB 7A1G[13]) is superposed to highlight its position near the mRNA entry channel. The figure was created using ChimeraX[45].

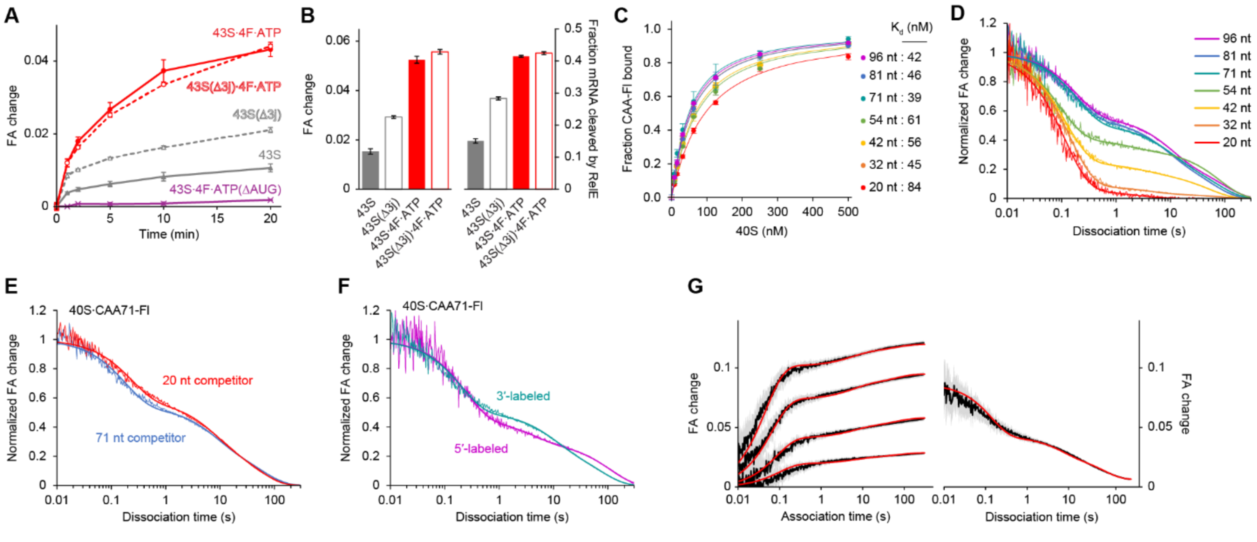

**Figure S2. Kinetic analysis of mRNA binding to the 40S subunit.**

**(A)** Start-site recognition kinetics monitored by FA changes of cap-CAA81(AUG)-FI under conditions as indicated. **(B)** Comparison of end-point FA changes vs. RelE cleavages (after 40 min incubation). **(C)** Apparent affinities of 3′-labeled CAA repeats for the 40S subunit measured by equilibrium FA changes. **(D)** 40S dissociation kinetics for 3′-labeled CAA RNAs of varying lengths. **(E)** 40S dissociation kinetics for CAA71-FI in the presence of excess unlabeled CAA71 or CAA20 competitor RNA. **(F)** 40S dissociation kinetics for CAA71 labeled at the 3′- or 5′-end. **(G)** Global fitting (red lines) for 40S·CAA71-FI association and dissociation kinetic dataset, based on the branched model. Gray areas represent σ intervals for each curve. All data are represented as mean ± SEM (n=3).

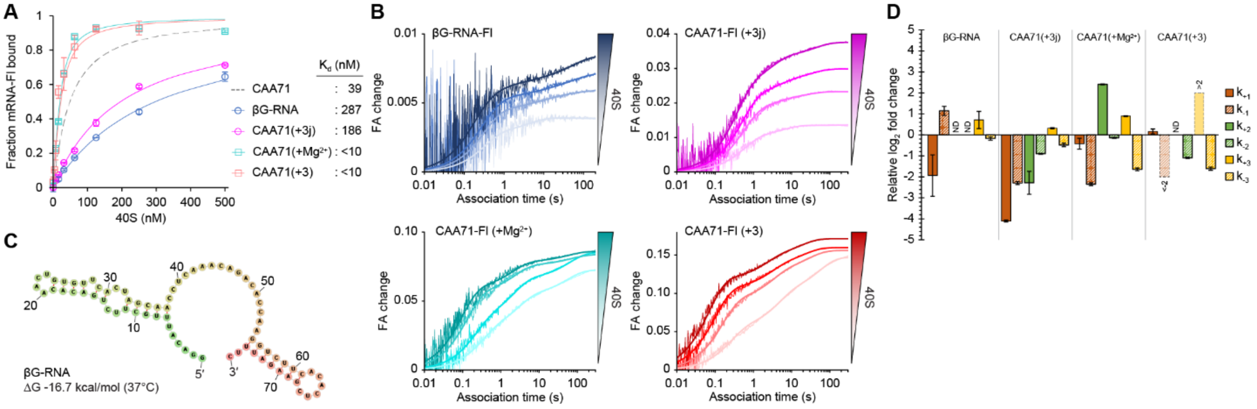

**Figure S3. Kinetic details under various conditions.**
**(A)** Apparent affinities of CAA71-Fl and βG-RNA-Fl for the 40S subunit under various conditions. **(B)** Association kinetics under various conditions. **(C)** Predicted secondary structure of βG-RNA using RNAfold web server[46]. **(D)** Relative $\log_2$ fold changes in kinetic rate constants under various conditions. 'ND' stands for not determined. Bars contoured with dashed lines represent estimates for a visual purpose. All data are represented as mean ± SEM (n=3).

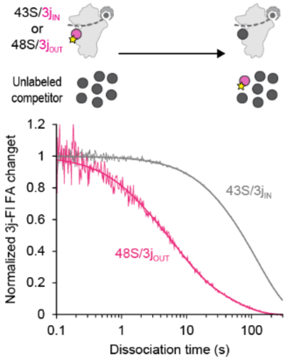


**Figure S4. eIF3j dissociation kinetics for 43S/3j$_{IN}$ and 48S/3j$_{OUT}$.**
Dissociation of 3j-Fl triggered by excess unlabeled competitor for the 43S (3j$_{IN}$ state) or the 48S (3j$_{OUT}$ state). Curves are fitted to double- or triple-exponential function as appropriate (thick lines). All data are represented as mean ± SEM (n=3).

**Table 1. Kinetic and thermodynamic parameters of CAA-Fl for the 40S binding**

| | CAA20 | CAA32 | CAA42 | CAA54 | CAA71 | CAA81 | CAA96 |
|---|---|---|---|---|---|---|---|
| $k_{-1}$ ($s^{-1}$) | 8.7 ± 0.5 | 7.0 ± 0.1 | 8.4 ± 0.4 | 10.5 ± 0.7 | 5.1 ± 0.2 | 5.6 ± 0.2 | 5.2 ± 0.4 |
| $k_{-2}$ ($s^{-1}$) | 0.45 ± 0.21 | 0.25 ± 0.03 | 0.34 ± 0.11 | 0.55 ± 0.11 | 0.11 ± 0.01 | 0.13 ± 0.01 | 0.11 ± 0.01 |
| $k_{-3}$ ($s^{-1}$) | ND | 0.012 ± 0.002 | 0.020 ± 0.001 | 0.012 ± 0.001 | 0.014 ± 0.001 | 0.017 ± 0.001 | 0.011 ± 0.001 |
| A1 | 0.94 ± 0.011 | 0.91 ± 0.002 | 0.75 ± 0.005 | 0.59 ± 0.003 | 0.49 ± 0.001 | 0.46 ± 0.005 | 0.44 ± 0.008 |
| A2 | 0.06 ± 0.011 | 0.05 ± 0.003 | 0.07 ± 0.005 | 0.08 ± 0.009 | 0.23 ± 0.002 | 0.26 ± 0.002 | 0.30 ± 0.005 |
| A3 | ND | 0.03 ± 0.001 | 0.18 ± 0.004 | 0.34 ± 0.006 | 0.27 ± 0.002 | 0.28 ± 0.004 | 0.26 ± 0.007 |
| $K_{d,app}$ (nM) | 84 ± 14 | 45 ± 12 | 56 ± 6 | 61 ± 2 | 39 ± 10 | 46 ± 5 | 42 ± 0.5 |

[*]Values are mean ± SEM (n=3). ND stands for Not Determined.

**Table 2. Kinetic and thermodynamic parameters of CAA71-Fl and βG-RNA-Fl for the 40S**

| | CAA71 | Global fitting[a] | βG-RNA | CAA71+$Mg^{2+}$ | CAA71+3j | CAA71+3 |
|---|---|---|---|---|---|---|
| $k_{-1}$ ($s^{-1}$)[b] | 5.1 ± 0.2 | 7.0 | 11.3 ± 2.4 | 1.0 ± 0.1 | 1.0 ± 0.1 | ND |
| $k_{-1}^{*}$ ($s^{-1}$)[c] | 6.1 ± 0.8 | | 2.8 ± 0.3 | 1.9 ± 0.5 | 1.5 ± 0.2 | <1 |
| $k_{-2}$ ($s^{-1}$)[b] | 0.11 ± 0.01 | 0.16 | ND | 0.10 ± 0.002 | 0.060 ± 0.001 | 0.052 ± 0.001[i] |
| $k_{-3}$ ($s^{-1}$)[b] | 0.014 ± 0.001 | 0.014 | 0.010 ± 0.001 | 0.004 ± 0.001 | 0.010 ± 0.001 | 0.005 ± 0.001 |
| $k_{+1}$ (μ$Ms^{-1}$)[c] | 80 ± 5 | 68 | 21 ± 12 | 60 ± 10 | 4.6 ± 0.1 | 89 ± 7 |
| $k_{+2}$ ($s^{-1}$)[d] | 0.053 ± 0.005 | 0.055 | ND | 0.279 ± 0.004 | 0.011 ± 0.003 | ND |
| $k_{+3}$ ($s^{-1}$)[d] | 0.0077 ± 0.0007 | 0.0071 | 0.027 ± 0.008 | 0.030 ± 0.0004 | 0.020 ± 0.001 | >0.04[i] |
| A1[e] | 0.49 ± 0.001 | 0.54 | 0.31 ± 0.13 | 0.10 ± 0.003 | 0.31 ± 0.01 | ≤0.11[i] |
| A2[e] | 0.23 ± 0.002 | 0.20 | ND | 0.26 ± 0.002 | 0.06 ± 0.02 | ≤0.11[i] |
| A3[e] | 0.27 ± 0.002 | 0.26 | 0.69 ± 0.13 | 0.64 ± 0.005 | 0.63 ± 0.03 | 0.89 ± 0.02 |
| $K_{d,app}$ (nM)[f] | 39 ± 10 | | 287 ± 13 | <10 | 186 ± 6 | <10 |
| $K_{d1obs}$ (nM)[g] | 79 | | 1202 | ND | 599 | ND |
| $K_{d1calc}$ (nM)[h] | 65 | | 538 | 17 ± 1 | 227 | <10 |

[a]Values in this column were obtained by the global fitting
[b]Rate determined by the dissociation analysis.
[c]Rate determined by the observed association rate plot.
[d]Rate calculated according to the branched model: $k_{+2} = K2*k_{-2}$, $k_{+3} = K3*k_{-3}$, where K2 = A2/A1 and K3 = A3/A1
[e]Amplitude determined by the dissociation analysis.
[f]Apparent equilibrium dissociation constant determined by the experimental saturation curve.
[g]Dissociation constant according to $K_{d1obs}=K_{dapp}*(1+K2+K3)$
[h]Dissociation constant according to $K_{d1calc}=k_{-1}/k_{+1}$
[i]Estimated based on the first phase in dissociation kinetics (see Methods)
[*]Values are mean ± SEM (n=3). ND stands for Not Determined.

**Table 3. Kinetic parameters of mRNA association with the 43S**

| | cap-CAA71 | | | | CAA71 |
|---|---|---|---|---|---|
| | 43S | 43S·4F·ATP | 43S·4F·ADP | 43S·4F·ATP(ΔE) | 43S·4F·ATP |
| $k_{1obs}$ ($s^{-1}$)[a] | 1.30 ± 0.13 | 1.97 ± 0.08 | 1.65 ± 0.13 | 2.13 ± 0.32 | 2.75 ± 0.57 |
| $k_{2obs}$ ($s^{-1}$)[a] | 0.22 ± 0.03 | 0.43 ± 0.02 | 0.28 ± 0.04 | 0.41 ± 0.09 | 0.51 ± 0.04 |
| $k_{3obs}$ ($s^{-1}$)[a] | 0.037 ± 0.004 | 0.073 ± 0.005 | 0.051 ± 0.007 | 0.060 ± 0.006 | 0.086 ± 0.010 |
| $A_{1obs}$[a] | 0.39 ± 0.02 | 0.38 ± 0.03 | 0.43 ± 0.04 | 0.49 ± 0.05 | 0.37 ± 0.04 |
| $A_{2obs}$[a] | 0.40 ± 0.01 | 0.54 ± 0.02 | 0.43 ± 0.04 | 0.46 ± 0.05 | 0.55 ± 0.02 |
| $A_{3obs}$[a] | 0.20 ± 0.01 | 0.08 ± 0.01 | 0.14 ± 0.02 | 0.05 ± 0.01 | 0.08 ± 0.01 |
| $A_{2obs}/A_{3obs}$[b] | 1.00 ± 0.02 | 3.51 ± 0.15 | 1.59 ± 0.23 | 4.61 ± 0.65 | 3.65 ± 0.46 |
| | cap-βG-RNA | | | | βG-RNA |
| | 43S | 43S·4F·ATP | 43S·4F·ADP | 43S·4F·ATP(ΔE) | 43S·4F·ATP |
| $k_{1obs}$ ($s^{-1}$)[a] | 0.39 ± 0.11 | 1.35 ± 0.15 | 0.66 ± 0.06 | 0.70 ± 0.06 | 1.25 ± 0.22 |
| $k_{2obs}$ ($s^{-1}$)[a] | 0.034 ± 0.004 | 0.127 ± 0.002 | 0.056 ± 0.002 | 0.049 ± 0.001 | 0.127 ± 0.005 |
| $k_{3obs}$ ($s^{-1}$)[a] | 0.0072 ± 0.0005 | 0.0093 ± 0.0005 | 0.0094 ± 0.0005 | 0.0083 ± 0.0004 | 0.0085 ± 0.0003 |
| $A_{1obs}$[a] | 0.12 ± 0.01 | 0.13 ± 0.01 | 0.18 ± 0.01 | 0.11 ± 0.01 | 0.17 ± 0.01 |
| $A_{2obs}$[a] | 0.49 ± 0.02 | 0.65 ± 0.01 | 0.44 ± 0.01 | 0.56 ± 0.02 | 0.59 ± 0.02 |
| $A_{3obs}$[a] | 0.39 ± 0.03 | 0.23 ± 0.02 | 0.39 ± 0.01 | 0.33 ± 0.01 | 0.24 ± 0.02 |
| $A_{2obs}/A_{3obs}$[b] | 1.00 ± 0.15 | 2.28 ± 0.25 | 0.88 ± 0.03 | 1.35 ± 0.09 | 1.95 ± 0.19 |

[a]Observed rate or amplitude in association analysis
[b]$A_{2obs}/A_{3obs}$ normalized to that for the 43S.
[*]Values are mean ± SEM (n=3).

### Table 4. Kinetic parameters of eIF3j-Fl dissociation

| Starting complex | 43S·4F·ATP | 43S·4F·ADP | 43S·4F·ATP(Δ4E) |
|---|---|---|---|
| Triggered by | cap-CAA71 | cap-CAA71 | cap-CAA71 |
| $k_1$ ($s^{-1}$) | 0.61 ± 0.03 | 0.61 ± 0.05 | 0.59 ± 0.01 |
| $k_2$ ($s^{-1}$) | 0.061 ± 0.003 | 0.045 ± 0.015 | 0.061 ± 0.006 |
| $A_1$ | 0.73 ± 0.01[a] | 0.15 ± 0.01[a] | 0.68 ± 0.01[a] |
| $A_2$ | 0.27 ± 0.01[a] | 0.09 ± 0.02[a] | 0.30 ± 0.01[a] |
| $\Delta r$[b] | 0.043 ± 0.001 | 0.010 ± 0.001 | 0.042 ± 0.001 |
| Starting complex | 43S·4F·ATP | 43S·4F·ADP | 43S·4F·ATP(Δ4E) |
| Triggered by | cap-GlobinNluc | cap-GlobinNluc | cap-GlobinNluc |
| $k_1$ ($s^{-1}$) | 0.79 ± 0.16 | 0.32 ± 0.03 | 0.21 ± 0.03 |
| $k_2$ ($s^{-1}$) | 0.0070 ± 0.0005 | 0.0048 ± 0.0014 | 0.0071 ± 0.0004 |
| $A_1$ | 0.50 ± 0.01[a] | 0.18 ± 0.03[a] | 0.19 ± 0.02[a] |
| $A_2$ | 0.50 ± 0.01[a] | 0.18 ± 0.02[a] | 0.73 ± 0.11[a] |
| $\Delta r$[b] | 0.013 ± 0.001 | 0.004 ± 0.001 | 0.012 ± 0.001 |
| Starting complex | 43S·4F cap-CAA71 | 43S | 43S·4F·ATP cap-CAA71 |
| Triggered by | ATP | Unlabeled eIF3j competitor | Unlabeled eIF3j competitor |
| $k_1$ ($s^{-1}$) | 0.79 ± 0.11 | 0.042 ± 0.004 | 0.73 ± 0.17 |
| $k_2$ ($s^{-1}$) | 0.28 ± 0.03 | 0.0075 ± 0.0003 | 0.13 ± 0.01 |
| $k_3$ ($s^{-1}$) | 0.022 ± 0.010 | | 0.020 ± 0.002 |
| $A_1$ | -0.30 ± 0.11[c] | 0.15 ± 0.02 | 0.26 ± 0.04 |
| $A_2$ | 0.82 ± 0.04 | 0.85 ± 0.02 | 0.49 ± 0.03 |
| $A_3$ | 0.18 ± 0.04 | | 0.26 ± 0.01 |
| $\Delta r$[b] | 0.025 ± 0.004 | 0.104 ± 0.009 | 0.038 ± 0.002 |

[a]Amplitude normalized to the total anisotropy change for 43S·4F·ATP in the presence of the corresponding mRNA.
[b]Total reduction in eIF3j-Fl anisotropy.
[c]Negative value stands an increase in anisotropy.
[*]Values are mean ± SEM (n=3).

### Table 5. List of mRNAs used in this study

| Name | Sequence |
|---|---|
| CAA20 | GGACAACAAAUGCAACAACA |
| CAA32 | GGACAACAACAACAAACCCUCGAACAACAACA |
| CAA42 | GGACAACAACAACAAACCCUCGAACAACAACAACAACAACAA |
| CAA54 | GGACAACAACAACAAACCAUGGAACAACAACAACAACAACAACAACAACAACAA |
| CAA71 | GGACAACAACAACAACAACAACAACAACAACAACAACAACAACAACAACAACAACAACA ACAACAACAACA |
| CAA81 | GGACAACAACAACAACAACAACAACAACAACAACAACAACAACAACAACAACAACAACA ACAACAACAACAACAACAACAA |
| CAA81(AUG) | GGACAACAACAACAAACCCUCGAACAACAACAACAACAACAACACCAUGGACAACC AUGGCAACAACAACAACAACAA |
| CAA96 | GGACAACAACAACAACAACAACAACAACAACAACAACAACAACAACAACAACAACAACA ACAACAACAAACCAUGGAACAACAACAACAACAACAA |
| βG-RNA | GGACAUUUGCUUCUGACACAACUGUGUUCACUAGCAACCUCAAACAGACACCAAGGUCU UCACACUCGAAGAUUUC |
